# EGF and IgA in maternal milk, donor milk and milk fortifiers in the Neonatal Intensive Care Unit setting

**DOI:** 10.1101/2024.10.26.620426

**Authors:** Christian Tamar, Kara Greenfield, Katya McDonald, Emily Levy, Jane Brumbaugh, Kathryn Knoop

**Author notes:** Correspondence to: Kathryn Knoop.

## Abstract

Human milk contains a variety of factors that positively contribute to neonatal health, including epidermal growth factor (EGF) and immunoglobulin A (IgA). When maternal milk cannot be the primary diet, maternal milk alternatives like donor human milk or formula can be provided. Donor human milk is increasingly provided to infants born preterm or low birth weight with the aim to supply immunological factors at similar concentrations to maternal milk. We sought to assess the concentrations of human EGF and IgA in the diet and stool of neonates between exclusive maternal milk, donor human milk, or formula-based diets. Using a prospective cohort study, we collected samples of diet and stool weekly from premature and low birth weight neonates starting at 10 days postnatal through five weeks of life while admitted to a neonatal intensive care unit (NICU). Compared to formula, there was significantly more EGF in both the milk and the stool of the infants fed human milk. Donor milk pooled from multiple donors contained similar concentrations of EGF and IgA to maternal milk, which was also significantly more compared to formula diets. Maternal milk fortified with fortifier derived from human milk contained significantly more EGF and IgA compared to unfortified maternal milk or maternal milk fortified with fortifier derived from bovine milk. Further analysis of human milk-derived fortifiers confirmed these fortifiers contained significant concentrations of EGF and IgA, contributing to an increased concentration of those factors when added to maternal milk compared to bovine milk-derived fortifiers. These findings illustrate how the choice of diet for a newborn, and even how that diet is modified through fortifiers or pasteurization before ingestion, impacts the beneficial biomolecules the infant receives from feeding.

## Introduction

Every year approximately 10% of infants in the United States are born prematurely at less than 37 weeks gestational age, putting the infant at risk for a myriad of developmental and chronic diseases [1]. Of prematurely born infants, those born at very low birth weight (VLBW), defined as less than 1500 grams [2], represent around 1% of all live births in the US [1]. These VLBW infants often spend weeks in the neonatal intensive care unit (NICU) as they develop and stabilize outside of the womb. During their stay in the NICU, VLBW infants are highly vulnerable to opportunistic and nosocomial pathogens. Maternal milk, defined here as milk received from the infants’ biological mother, is recommended as the exclusive diet for infants during the critical first six months of life by the American Academy of Pediatrics and World Health Organization [3]. Early enteral feeding with maternal milk reduces incidence of, and mortality from, infectious diseases, including outcomes like neonatal late-onset sepsis [4] and necrotizing enterocolitis [5, 6].

Many factors in human milk confer neonatal immunity and temper inflammatory responses to unfamiliar antigens, protecting against infection and shaping the infant microbiome [7-9]. One of these human milk factors, epidermal growth factor (EGF), has been shown to improve intestinal barrier function by promoting epithelial cell growth and decreasing bacterial translocation in maternal milk-fed infants [10-13]. Furthermore, in a mouse model of neonatal sepsis, EGF improved intestinal barrier function in neonates, prevented enteric pathogens translocating from the intestine, and prevented the development of sepsis secondary to bloodstream infection [14]. While EGF directly strengthens the intestinal epithelium, the human milk factor immunoglobulin A (IgA) contributes to neonatal intestinal health primarily through its interactions with the developing microbiome. In addition to encouraging the growth of bacterial commensals, maternal IgA provides passive immunity to the neonate through targeted protection against immunologically relevant antigens before the infant gains the ability to generate their own IgA response, reducing the risk of enteric infection and necrotizing enterocolitis [15-18]

When the maternal milk containing these valuable factors is unavailable, Donor human milk and human milk-derived fortifiers are increasingly utilized for VLBW infants rather than formula [19, 20]. Donor milk can be expressed throughout a donor’s lactation cycle, often months following parturition when the concentrations of many milk factors, including EGF and IgA, are lower than immediately after birth [21-24]. We have observed in a preclinical model that milk expressed closer to parturition contained more EGF and offered more protection from enteric pathogens than milk expressed later in lactation [14]. This gradual decline throughout lactation is reflected in the stool of infants fed maternal milk but not in formula-fed infants [14]. Donor milk is often provided in this asynchronous manner, with the timing of donor milk collection during lactation unlikely to be matched to the infant’s corrected gestational age. To address how much the composition of human milk can naturally vary between donors and minimize the impact of individual sample differences, some milk banks pool their donor milk between multiple donors [25]. In addition to the choice of primary diet between maternal milk, donor milk, and formula, a further measure to address the high nutritional demands of preterm or VLBW infants is the introduction of a fortifier to their milk. Fortifiers can be derived from different origins, including human and bovine milk, and can vary substantially in their nutritional composition, with potential ramifications for their immunological benefit [26].

In this study, we analyzed the concentrations of EGF and IgA in ready-to-feed maternal milk, donor milk, and formula provided to neonates. In addition to these diets, we measured EGF in the stool of infants to determine if the amount of EGF present in the stool reflected the amount received in the diet. Finally, we examined EGF and IgA concentrations in different types of human milk fortifiers. Here, we show that EGF and IgA concentrations vary between maternal milk alternative diets, including between individual and pooled donor milk, and can be further altered by the addition of different types of human milk fortifiers, with potential functional consequences.

## Methods

### Human Subjects

Mother-infant paired participants were recruited for this observational study between March 2021-March 2023 based on infant birth weight (<2500 grams) or pre-term birth at <36 weeks gestation. Infant participants were at least 3 days old at enrollment and admitted to the Mayo Clinic NICU, in Rochester, Minnesota, USA. Beginning at 8 days following birth or later, milk and infant stool were collected weekly for 5 weeks while hospitalized. Exclusion criteria included major congenital anomalies involving the intestinal tract and placement on nil per os (NPO) orders. Written informed consent was obtained from a parent of eligible infants. Clinical data including gestational age at birth, birth weight, birth length, frontal occipital circumference, mode of delivery, age at full feeding, history of culture positive sepsis, use of steroids or antibiotics, and diet including fortification was collected and deidentified. This study was approved through the Mayo Clinic Institutional Review Board.

### Milk and Stool Collection

A 2 mL aliquot of the diet for the participant was saved from the ‘ready-for-feed’ milk by the nursing staff immediately prior to feeding and stored at 4°C until collection for processing for no more than 4 hours. Between 24 and 48 hours later, stool was collected from the diaper by nursing staff using a sporked collection tube and stored at -20°C until collection for processing within 4 hours. Incomplete sample pairs were discarded. Diets consisted of expressed maternal milk, donor milk, or infant formula, as available or based on the discretion of the clinical care team. Maternal milk, following expression, was stored at 4°C until utilized for feed. Donor milk was obtained from the Ohio Mothers’ Milk Bank, which pools donor milk from 3-5 individuals, and batch information was collected. When fortified, diet was fortified with bovine-based fortifiers (Similac Human Milk Fortifier or Enfamil Human Milk Fortifier) or human milk-derived fortifiers (Prolacta +4, +6, +8), at the discretion of the clinical care team. Milk type and fortifiers were noted by staff at time of collection.

### Individual Human Donor Milk

In some experiments, individual donor milk was used for experimental assays. Individual donor milk was obtained from the Minnesota Milk Bank for Babies. Pasteurized milk frozen at -80°C from 20 different individual donors was obtained and kept separate. Time of collection post-partum was collected.

### Fortifiers and Formula

Some participants’ diets were supplemented with either bovine milk-based fortifiers (Similac Human Milk Fortifier or Enfamil Human Milk Fortifier) or human milk-based fortifiers (Prolact+6 H^2^MF, Prolact+8 H^2^MF Human Milk Fortifier), at the discretion of the clinical care team. In some experiments, fortifiers were used for experimental assays and aliquots of fortifiers were obtained for direct analysis from the Mayo Clinic Nutrition Lab. For direct analysis of fortifiers, independent lots were used as biological replicates.

### ELISA

Human breast milk samples, human milk fortifiers and human stool samples were analyzed by enzyme-linked immunoabsorbent assay (ELISA) for human IgA (88-50600, Invitrogen, Waltham, MA, USA) and human EGF (DY236, R&D Systems, Minneapolis, MN, USA), according to the manufacturer’s protocol. Milk samples analyzed for human IgA content were diluted from 1:1000 to 1:20,000, samples analyzed for human EGF content were diluted from 1:500 to 1:2500, and human milk fortifiers were diluted at 1:10. Samples were diluted with a 1:20 dilution of Assay Buffer A Concentrate (PBS with 1% Tween -20 and 10% BSA, Invitrogen 88-50600-88) in deionized water. For stool sample preparation, 100 mg of stool was taken from the sample and homogenized in 1 mL of phosphate buffered solution (PBS). Sample optical density was found using a BioTek 800 TS absorbance reader (BioTek, Winooski, VT, USA) at 450nm and 570nm. Tests for human IgA bacterial binding began by coating a 96-well ELISA plate with either heat-killed *Streptococcus agalactiae* (Group B Strep) or heat-killed *Escherichia coli*, then subsequently adhered to the manufacturer’s protocol (Invitrogen 88-50600).

### Statistics

All statistical analysis was done using GraphPad Prism 9.0 (GraphPad Software Inc., Boston, MA, USA). Data is reported as mean +/-the standard error of the mean. Differences between groups were analyzed using two-way ANOVAs, with Sidak’s posttest, Kruskal-Wallis test with Dunn’s multiple comparisons, and Pearson’s Correlation.

## Results

### EGF is decreased in individual donor milk, but not pooled donor milk

Our enrolled cohort consisted of 74 mother-infant dyads, with 237 total diet samples collected which were recorded as either maternal milk (218), donor milk (13), or formula (6). 243 stool samples were also collected from the same infants. 20 samples of pasteurized milk from individual donors were obtained from the Minnesota Milk Bank for Babies for further comparison. Additional clinical characteristics of the cohort are displayed in Table 1. Relative to maternal milk (59.72 ± 2.25 ng/mL), both formula (9.54 ± 1.48 ng/mL) and individual donor milk samples (23.74 ± 2.29 ng/mL) had significantly decreased EGF (Fig. 1A). Pooled donor milk samples (70.15 ± 7.14 ng/mL) contained similar EGF concentrations as compared to maternal milk samples (Fig. 1A). To determine if the amount of EGF in the stool of infants reflects the initial amount of EGF from the diet, stool was collected from the infants receiving these milk or formula samples and evaluated for EGF content (Fig. 1B). Infants fed maternal milk (248.33 ± 40.62 pg/g) or pooled donor milk (194.59 ± 51.85 pg/g) had significantly more EGF in their stool on average than those given formula (29.06 ± 5.47 pg/g). We observed a positive correlation when plotting the individual infants’ stool EGF against the EGF contents of paired milk or formula samples (Fig. 1C).

**Table 1.**
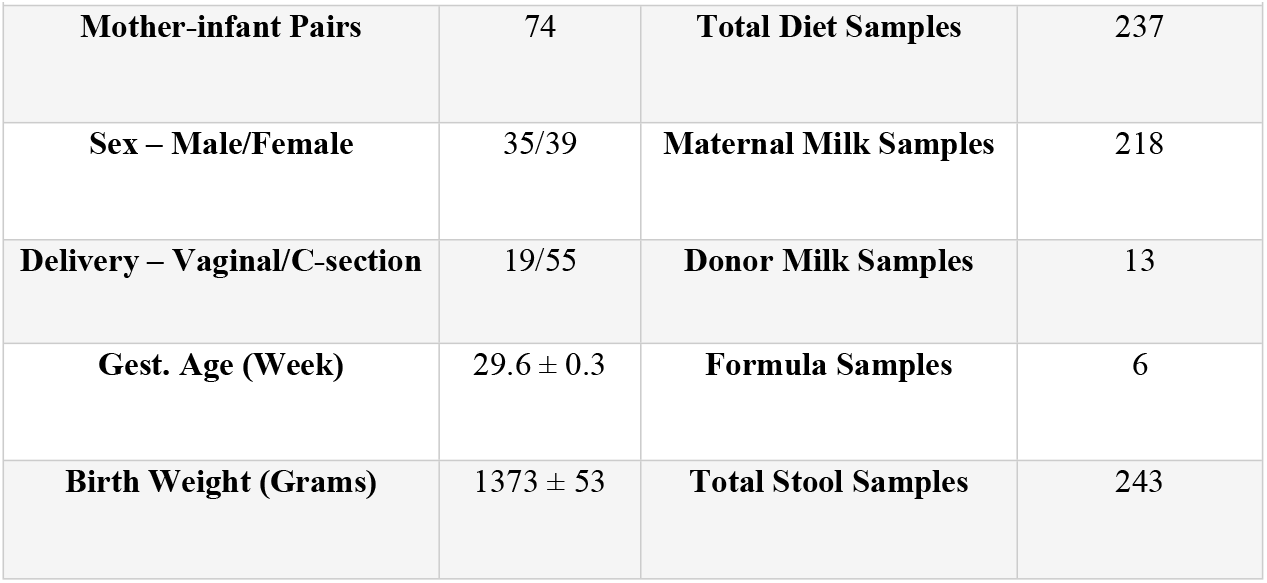
Description of Clinical Cohort.

**Fig. 1:**
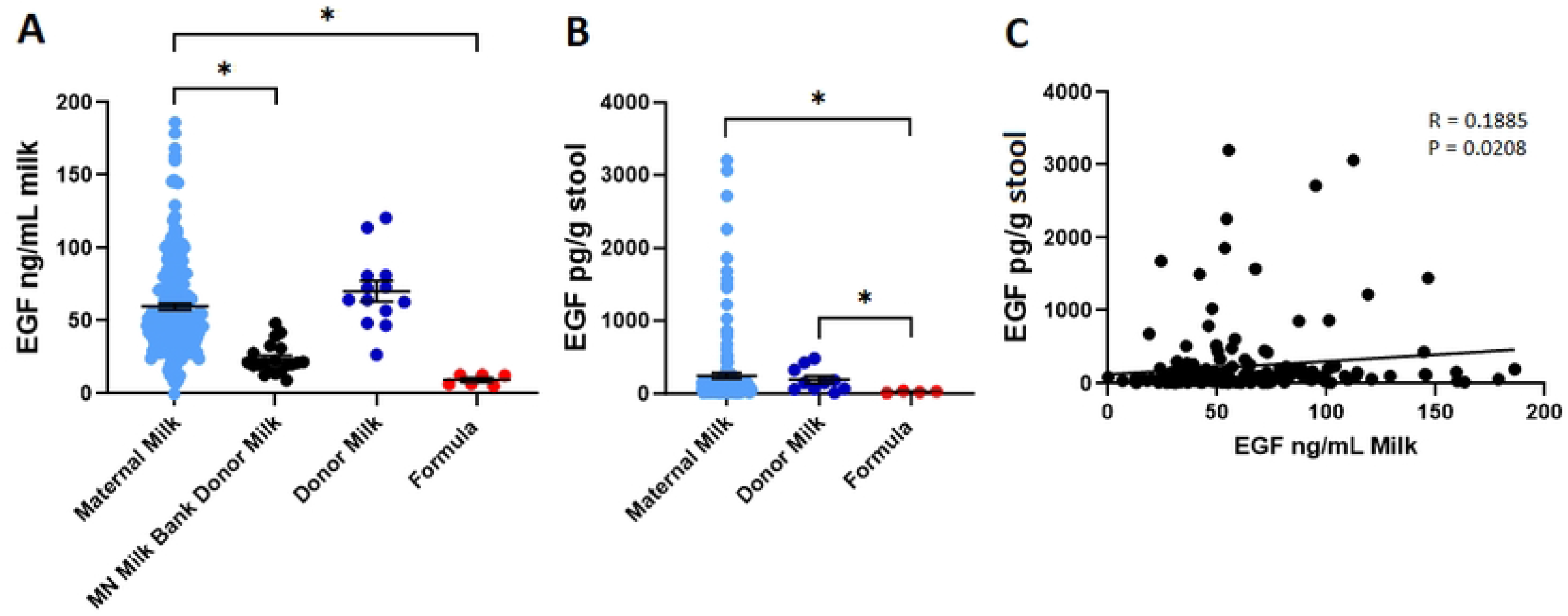
EGF is decreased in individual donor milk, but not pooled donor milk. A) Concentration of EGF in maternal milk (light blue, n=218), individual donor milk (black, n=20), pooled donor milk (dark blue, n=13), and formula (red, n=6). B) Concentration of EGF in stool from infants fed maternal milk (light blue, n=163), donor milk (dark blue, n=10), or formula (red, n=4). (C) EGF in stool plotted against EGF in milk from matched-pairs specimens. * denotes significance, p<0.05, Kruskal-Wallis test with Dunn’s multiple comparisons test in A and B, Pearson’s correlation in C.

### Human milk-based fortifiers contribute to EGF concentration

We next assessed if fortifiers affected the concentration of EGF by comparing the fortified or unfortified milk (Fig. 2A). Fortified maternal milk (65.29 ± 2.86 ng/mL) had significantly increased EGF compared to unfortified maternal milk (49.11 ± 3.28 ng/mL), though this was not the case in fortified donor milk as compared to unfortified donor milk. The significant differences of EGF in the milk were not detected in infant stool (Fig. 2B). Following assessment based on type of fortifier used (bovine milk-derived or human milk-derived), we observed milk fortified with a human milk-derived fortifier contained significantly more EGF (74.16 ± 3.43 ng/mL) as compared to unfortified maternal milk (49.11 ± 3.31 ng/mL), or milk fortified with a bovine milk-derived fortifier (41.29 ± 2.57 ng/mL; Fig. 2C). Measuring fortifiers directly, only human milk-derived fortifier products exhibited a substantial level of human EGF (Fig. 2D).

**Fig. 2:**
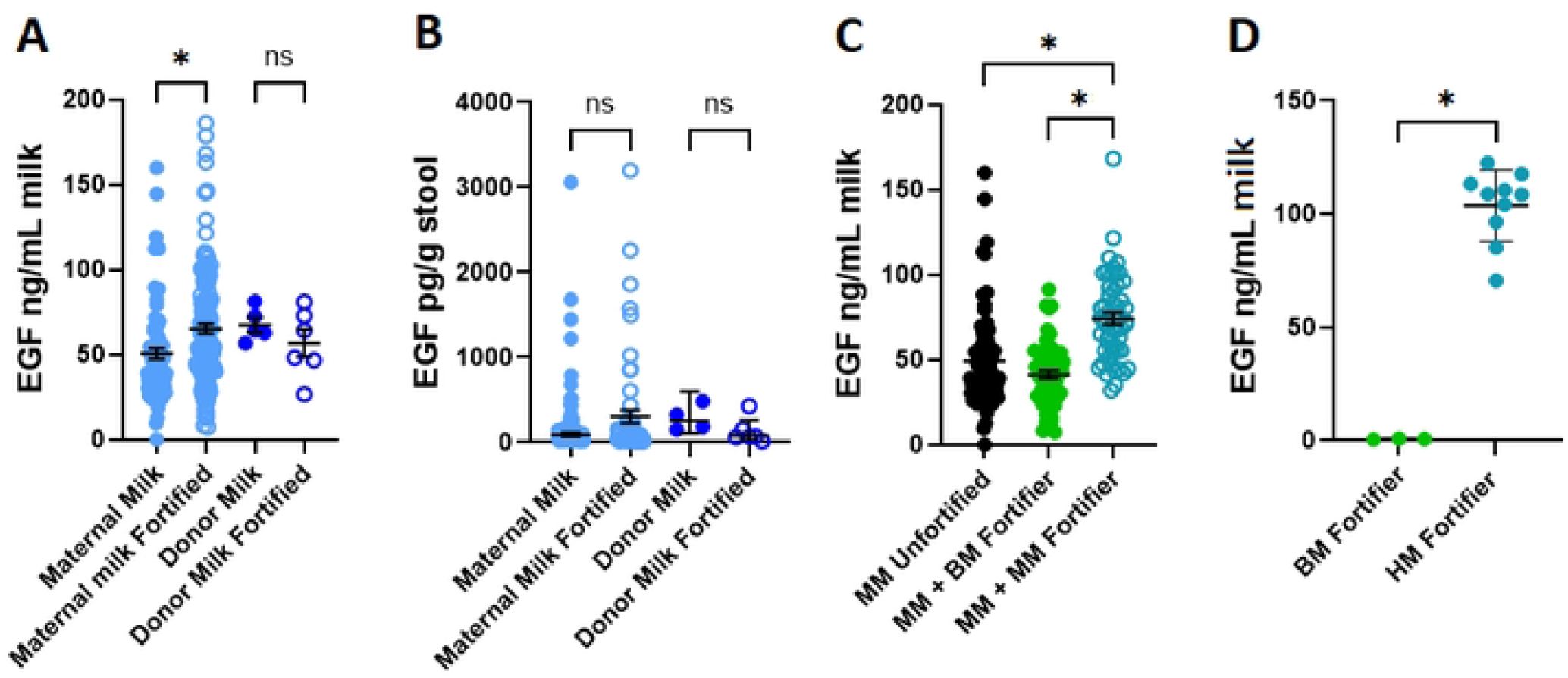
Human milk-based fortifiers contribute to EGF concentration. A) Concentration of EGF in maternal milk (light blue) or donor milk (dark blue), unfortified (solid circles) or fortified (open circles) (maternal milk unfortified: n=75; maternal milk fortified: n=143; donor milk unfortified: n=5; donor milk fortified: n=6). B) Concentration of EGF in stool from infants’ maternal milk (light blue) or donor milk (dark blue), unfortified (solid circles) or fortified (open circles) (maternal milk unfortified: n=90; maternal milk fortified: n=62; donor milk unfortified: n=4; donor milk fortified: n=6). C) Concentration of EGF in unfortified maternal milk (black, n=75), maternal milk fortified with bovine milk-based fortifier (green, n=55), or maternal milk fortified with human milk-based fortifier (blue, n=45). D) Concentration of EGF in bovine milk-based fortifier (green) and human milk-based fortifier (blue). * denotes significance, p<0.05 Kruskal-Wallis test with Dunn’s multiple comparisons test in A, B, C and D.

### IgA is decreased in individual donor milk and is present in human milk-based fortifiers

We then evaluated the potential variation of IgA in maternal and donor milk. As with EGF, maternal milk (346.35 ± 17.91 ug/mL) and pooled donor milk samples (327.86 ± 93.01 ug/mL) had substantially higher IgA concentrations compared to individual donor milk (93.64 ± 12.78 ug/mL) and formula samples (18.76 ± 4.05 ug/mL; Fig. 3A). We observed a positive correlation between the IgA concentrations of milk samples and those samples’ corresponding EGF concentrations (Fig. 3B). We found increased IgA in maternal milk with a human milk-based fortifier (445.93 ± 44.53 ug/mL) compared to unfortified maternal milk (301.8 ± 24.77 ug/mL), which was not observed in maternal milk fortified with a bovine milk-based fortifier (304.75 ± 32.37 ug/mL; Fig. 3C). Furthermore, when the fortifiers alone were evaluated for human IgA, human milk-based fortifiers contained significantly more IgA than the bovine milk-based fortifier, which contained negligible human IgA (Fig. 3D).

**Fig. 3:**
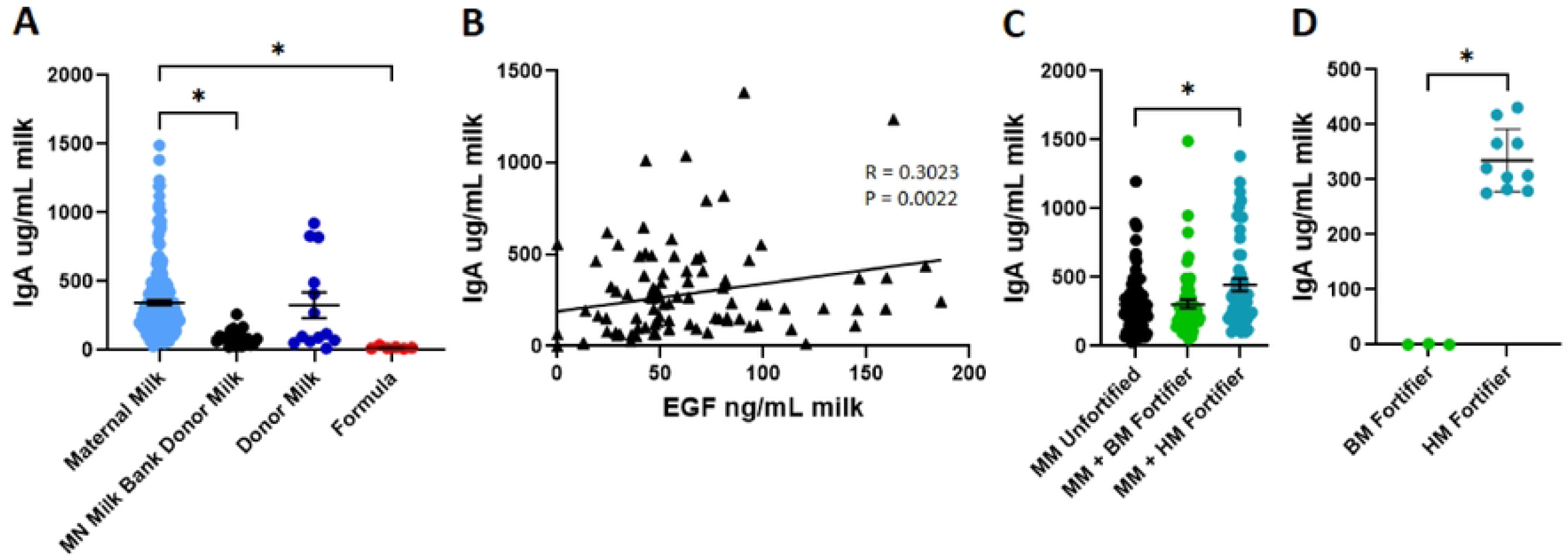
IgA is decreased in individual donor milk and is present in human milk-based fortifiers. A) Concentration of IgA in maternal milk (light blue, n=216), individual donor milk specimens (black, n=20), pooled donor milk specimens (dark blue, n=13) and formula (red, n=6). B) IgA in milk plotted against EGF in milk from same specimen. C) Concentration of IgA in unfortified maternal milk (black, n=75), maternal milk fortified with bovine milk-based fortifier (green, n=55) and maternal milk fortified with human milk-based fortifier (blue, n=52). D) Concentration of IgA in bovine milk-based fortifier (green) and human milk-based fortifier (blue). * denotes significance, p<0.05, Kruskal-Wallis test with Dunn’s multiple comparisons test in A, C, and D, Pearson’s correlation in B.

### IgA retained in human milk-based fortifiers is cross-reactive to potential pathogens

To determine if IgA present in the human milk-based fortifiers retained functional activity, we tested the ability of IgA to react and bind common pathogenic bacteria *Streptococcus agalactiae* (also known as Group B *Streptococcus*, or GBS) and *Escherichia coli* by ELISA. We detected significant binding of IgA from human milk-based fortifiers to both GBS and *E. coli* (Fig. 4A-B). Finally, we assessed binding of fortified and unfortified maternal milk to GBS and *E. coli*. While we did not observe a statistically significant increase in absorbance from the human milk-based fortified milk compared to the bovine milk-based fortified milk or unfortified milk with either GBS or E. coli (Fig. 4C-D), we did observe a trend of the bovine milk-based fortified milk samples displaying less IgA binding than the other two groups, though the trend was not statistically significant.

**Fig. 4:**
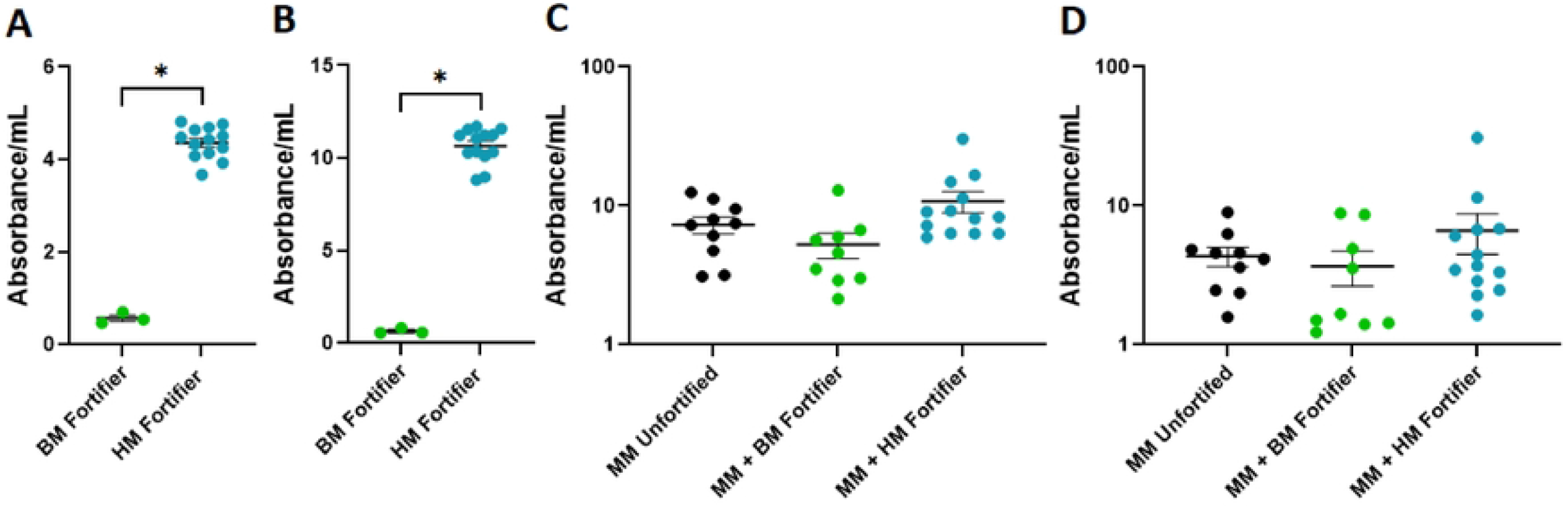
IgA retained in human milk-based fortifiers is cross-reactive to potential pathogens. A) Reactivity of IgA in bovine milk-based fortifier (green) and human milk-based fortifier (blue) to *Streptococcus agalactiae* (GBS). B) Reactivity of IgA in bovine milk-based fortifier (green) and human milk-based fortifier (blue) to *Escherichia coli*. C) Reactivity of IgA in unfortified maternal milk (black), maternal milk fortified with bovine milk-based fortifier (green) and maternal milk fortified with human milk-based fortifier (blue) to *Streptococcus agalactiae* (GBS). D) Reactivity of IgA in unfortified maternal milk (black), maternal milk fortified with bovine milk-based fortifier (green) and maternal milk fortified with human milk-based fortifier (blue) to *Escherichia coli*. * denotes significance, p<0.05, Kruskal-Wallis test with Dunn’s multiple comparisons test in A, B, C and D.

## Discussion

Our findings demonstrate that EGF and IgA content delivered to infants in the NICU can vary substantially depending on the diet type they receive and on further dietary additions like human milk fortifiers. Concerning diet choice, our results indicate a pooled donor milk diet may be more likely to provide greater EGF and IgA concentrations than individual donor milk or formula and would more closely reflect the concentrations of these factors delivered by a maternal milk diet. In addition, we found that a human milk-based fortifiers retained significant levels of these biomolecules and contributed to an increase in EGF and IgA when added to milk. Together, these findings further underline the importance of dietary choices in early life as a source of biologically functional human milk factors.

When we analyzed donor milk specimens from individual donors, EGF and IgA were both significantly reduced compared to maternal milk specimens, while pooled milk samples were similar to maternal milk. This is consistent with results from Young et al. (2020) [27] reporting that single donor pools contained significantly less IgA than multi-donor milk pools. Interestingly, we observed in our dataset that IgA concentrations in pooled donor milk mirrored those in maternal milk despite undergoing Holder Pasteurization, which has been shown to dramatically reduce IgA in treated milk [28], possibly as a result of the relatively small amount of pooled donor milk samples analyzed. One possible explanation for the difference in average EGF and IgA concentration between pooled donor milk and the individual donor milk samples that compose a pooled milk sample could be the presence of a small amount of high expressors in the donor pool. Milk component concentrations are highly variable between individuals [29] and can further vary based on prematurity [30], size at gestational age [31], and postpartum milk age [32], though Young et al. (2020) [27] found this to be a negligible factor for IgA specifically. Together, these contributions could lead to some donor milk samples being much more highly concentrated in certain milk factors than others, depending on the individual donor, where the inclusion of one of these samples in a donor milk pool would significantly increase the concentration of some milk factors in the final pooled sample, potentially bringing it more in line with average maternal milk. Our observed correlation between EGF and IgA concentrations in a given milk sample suggests that inclusion of individual donor milk highly concentrated in one milk factor within a donor pool may increase the concentration of other beneficial factors in the final pooled donor milk. However, a diet of individual donor milk still delivers more of these crucial biomolecules than a formula diet, which contains virtually no EGF or IgA. The capacity of pooled donor milk in replicating the EGF and IgA concentrations of maternal milk demonstrates that donor milk pooling could provide an immunological value that formula generally cannot, though larger cohorts of pooled donor milk-fed and formula-fed infants are needed to validate this trend.

In addition to the macronutrients and minerals that human milk fortifiers aim to supplement, we observed human milk-based fortifiers contain meaningful amounts of human EGF and IgA. Human milk-based fortifiers, unlike the bovine milk-based fortifier we analyzed, preserved enough of both biomolecules that fortified milk was significantly enriched for EGF and IgA compared to unfortified maternal milk. Previous work has observed differences in milk factor concentrations between milk fortified with either human or bovine derived fortifiers [33], though EGF has not been analyzed in this context and effects on IgA are inconclusive [34, 35]. We also found that the human IgA present in the human milk-based fortifiers was able to bind to multiple pathogens implicated in neonatal sepsis, suggesting that IgA present in human milk-derived fortifiers could be protective against sepsis events resulting from translocation of pathogens in the infant gut. This IgA may also play a role in shaping the composition of the developing microbiome [36], though further analysis and understanding of phenotypic differences associated with different dietary options is needed [37, 38]. While the impact of fortifier derivation from human or bovine sources is not clearly established [39-42] and the recommendations for milk fortification for preterm infants in general are still being determined [43-45], the retention of beneficial components like EGF and IgA in human milk-based fortifiers could be considered for a diet that aims to retain human milk-derived immunological protection.

## Limitations

One limitation of this study is a lack of association between dietary options and health outcomes like neonatal sepsis. Our study was unable to directly judge the health impacts of the various diets given its observational nature, as well as the rarity of neonatal infection outcomes. While our data concurs with other work reinforcing the value of pooling donor milk as a method to reduce individual variance between donor milk samples [27, 29, 32], further research is necessary to assess how the retention of critical factors in donor milk and fortifiers may contribute to the intestinal health of the infant. We also had significantly fewer donor milk samples than maternal milk samples in our dataset and were therefore unable to assess the contribution of different types of fortification to milk factor concentrations in donor milk. A further limitation was our inability to ask if reduced EGF in individual donor milk was reflected in the stool of corresponding infants, as this study did not have enrolled neonates on an individual donor milk diet due to current nutritional practices at the study’s site location. We did, however, find a significant amount of EGF in the stool of infants fed human milk (maternal or pooled donor) as compared to formula, which suggests that infants ingesting greater levels of EGF are also passing that EGF through their digestive tract. How this milk EGF is utilized biologically by the neonate through the gastrointestinal tract as a growth factor remains an active area of interest. One possible option to further investigate the functional significance of increased EGF and IgA would have been to analyze the stool microbiome compositions of our samples and relate the results to the amount of EGF and IgA in each infants’ diet. Finally, as our primary focus was neonates’ exposure to milk factors from ready-to-feed diets, we did not account for potential influences from processing factors upstream of diet delivery.

## Acknowledgements

Author contributions: JB, EL and KK conceived the study. CT, KG, and KM processed specimens, ran ELISAs, and analyzed data. CT and KK wrote the initial manuscript, all authors critically reviewed and approved the manuscript.

